# A LaeA- and BrlA-dependent cellular network governs tissue-specific secondary metabolism in the human pathogen *Aspergillus fumigatus*

**DOI:** 10.1101/196600

**Authors:** Abigail L. Lind, Fang Yun Lim, Alexandra A. Soukup, Nancy P. Keller, Antonis Rokas

## Abstract

Biosynthesis of many ecologically important secondary metabolites (SMs) in filamentous fungi is controlled by several global transcriptional regulators, like the chromatin modifier LaeA, and tied to both development and vegetative growth. In *Aspergillus* molds, asexual development is regulated by the BrlA>AbaA>WetA transcriptional cascade. To elucidate BrlA pathway involvement in SM regulation, we examined the transcriptional and metabolic profiles of Δ*brlA*, Δ*abaA*, Δ*wetA* and wild-type strains of the human pathogen *Aspergillus fumigatus*. We find that BrlA, in addition to regulating production of developmental SMs, regulates vegetative SMs and the SrbA-regulated hypoxia stress response in a concordant fashion to LaeA. We further show that the transcriptional and metabolic equivalence of Δ*brlA* and Δ*laeA* is mediated by a LaeA requirement preventing heterochromatic marks in the *brlA* promoter. These results provide a framework for the cellular network regulating not only fungal SMs but diverse cellular processes linked to virulence of this pathogen.

## Introduction

Filamentous fungi produce a remarkable diversity of specialized secondary metabolites (SMs), which are small molecules that play diverse ecological roles in fungal defense, communication, and virulence (1). In fungi, SMs are typically produced by pathways organized into contiguous biosynthetic gene clusters (BGCs), an organization atypical of metabolic pathways in most other eukaryotes (2). The transcription of these BGCs is often controlled by both cluster-specific transcription factors as well as globally-acting transcriptional regulators. These global regulators respond to a variety of environmental signals including pH, temperature, light, and nutrient sources to transcriptionally regulate BGCs, and are typically well conserved in filamentous fungi (3).

Many of the environmental signals that regulate SM production in *Aspergillus* fungi, including temperature, pH, and carbon or nitrogen sources, also trigger the onset of asexual and sexual development (4). At the cellular level, this coupling between SM production and development is orchestrated in part by the velvet protein complex, which is composed of two velvet domain proteins, VeA and VelB, and the methyltransferase LaeA (5). Although the precise mechanism by which the velvet complex regulates the two processes is unknown, LaeA regulates transcription epigenetically through heterochromatin reorganization of target DNA (6, 7). The result of this coupling of SM and development is that several SMs show tissue specificity, i.e., they are localized or produced only in certain tissues. For example, the SMs DHN melanin, fumigaclavines, endocrocin, trypacidin, and fumiquinazolines, appear to be specifically produced in the asexual spores of *Aspergillus fumigatus* (8–12). Importantly, several of these asexual spore (conidial) metabolites, all LaeA-regulated (13), are part of the pathogenic arsenal of this human pathogen (reviewed in 14).

In *Aspergillus,* asexual development is controlled by three regulatory genes sequentially expressed at specific stages of asexual fruiting body development (conidiation) (14). The first protein of this regulatory cascade, BrlA, accumulates in vegetative cells shortly before asexual development (15). In the middle stages of conidiation, BrlA activates AbaA, which controls the development of the asexual fruiting body (conidiophore) and spores (conidia). In late stages of asexual development AbaA activates WetA, which is required for conidial maturation through governance of critical conidial cell wall components (15). Recent evidence suggests that these three regulators may also be involved in regulating the expression of BGCs whose SM products are specifically found in asexual spores (8, 12, 16, 17).

To shed light on the relationship between regulation of asexual development and tissue-specific regulation of secondary metabolism in filamentous fungi, we performed global transcriptomic and metabolomic analyses on Δ*brlA*, Δ*abaA*, Δ*wetA* and wild-type strains of *A. fumigatus* in conditions previously shown to induce the production of both vegetative growth-specific and asexual development-specific SMs (13). Our results show that BrlA positively regulates the transcriptional activity of 13 BGCs and their SMs; importantly, BrlA regulates not only the production of both asexual development-specific SMs and of vegetative growth-specific SMs, but also the activity of several transcriptional regulators of diverse cellular processes, such as the SrbA-regulated hypoxia stress response. Remarkably, comparison of BrlA-regulated and LaeA-regulated BGCs shows that nine BGCs (DHN melanin, fumigaclavine, endocrocin, trypacidin, helvolic acid, fumisoquin, gliotoxin, fumiquinazoline, and pyripyropene A) appear to be identically regulated by both proteins. To further dissect the regulatory overlap between LaeA and BrlA, we used ChIP-qPCR to show that *laeA* loss results in heterochromatic marks in the *brlA* promoter and hence dampening of *brlA* expression. The epigenetic and epistatic effect of *laeA* on *brlA* explains, to a large degree, the concordance of BGC regulation and hypoxia gene regulation by these two proteins. These results argue that LaeA and BrlA are key components of the cellular network governing tissue-specific secondary metabolism as well as of diverse cellular processes in filamentous fungi.

## Results and Discussion

### Genome-wide transcriptional impact of BrlA, AbaA, and WetA

To examine the genome-wide regulatory roles of the three central regulators of asexual development, we performed RNA sequencing on *A. fumigatus* wild-type (WT), Δ*brlA*, Δ*abaA*, and Δ*wetA* mutant strains grown on minimal media in conditions known to induce the production of both vegetative growth-specific and asexual development-specific SMs. 6,738 of the 9,784 genes in the genome of the *A. fumigatus* Af293 strain were differentially expressed in the Δ*brlA* versus WT comparison (3,358 over-expressed and 3,380 under-expressed) (**Table 1, Table S1**). Fewer genes were differentially expressed in the Δ*abaA* versus WT comparison (1,895 differentially expressed genes; 1,148 over-expressed and 747 under-expressed) and in the Δ*wetA* versus WT comparison (2,158 differentially expressed genes; 1,192 over-expressed and 966 under-expressed).

**Table 1.**
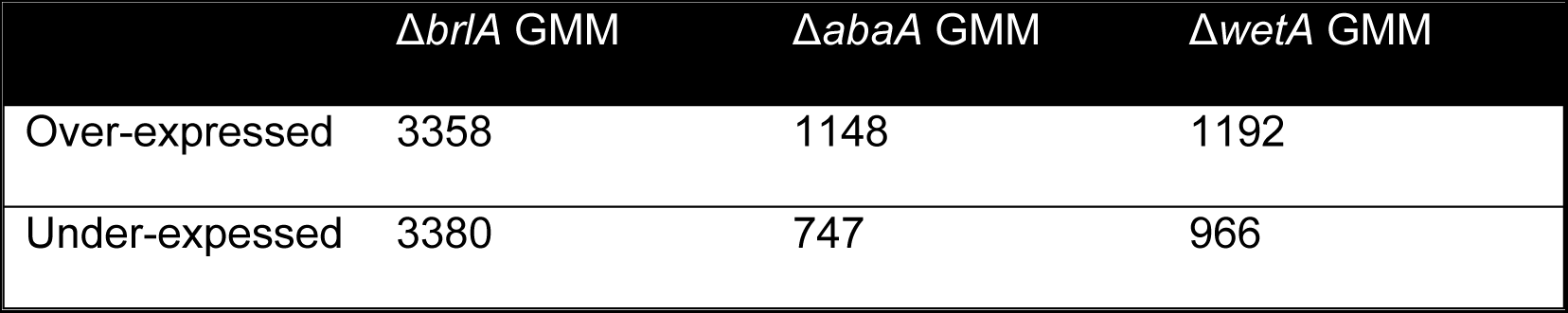
Differential gene expression in all RNA-seq comparisons.

| | $\Delta brlA$ GMM | $\Delta abaA$ GMM | $\Delta wetA$ GMM |
| --- | --- | --- | --- |
| Over-expressed | 3358 | 1148 | 1192 |
| Under-expressed | 3380 | 747 | 966 |

To determine processes positively regulated by BrlA, AbaA, and WetA, we performed GO enrichment analysis for genes under-expressed in each deletion strain. Among the 55 functional categories enriched in genes under-expressed in Δ*brlA* versus WT were secondary metabolism, response to stress, developmental process, asexual sporulation, cellular respiration, ribosome, mitochondrion, and structural molecule activity (**Figure 1A, Table S2**). Genes under-expressed in the Δ*abaA* versus WT were enriched for six functional categories, namely secondary metabolic process, oxidoreductase activity, cellular amino acid metabolic process, response to chemical stimulus, toxin metabolic process, and transferase activity (**Figure 1B, Table S2**). Finally, genes under-expressed in Δ*wetA* versus WT were enriched for three categories, which were secondary metabolic process, toxin metabolic process, and oxidoreductase activity (**Figure 1C, Table S2**).

**Figure 1.**
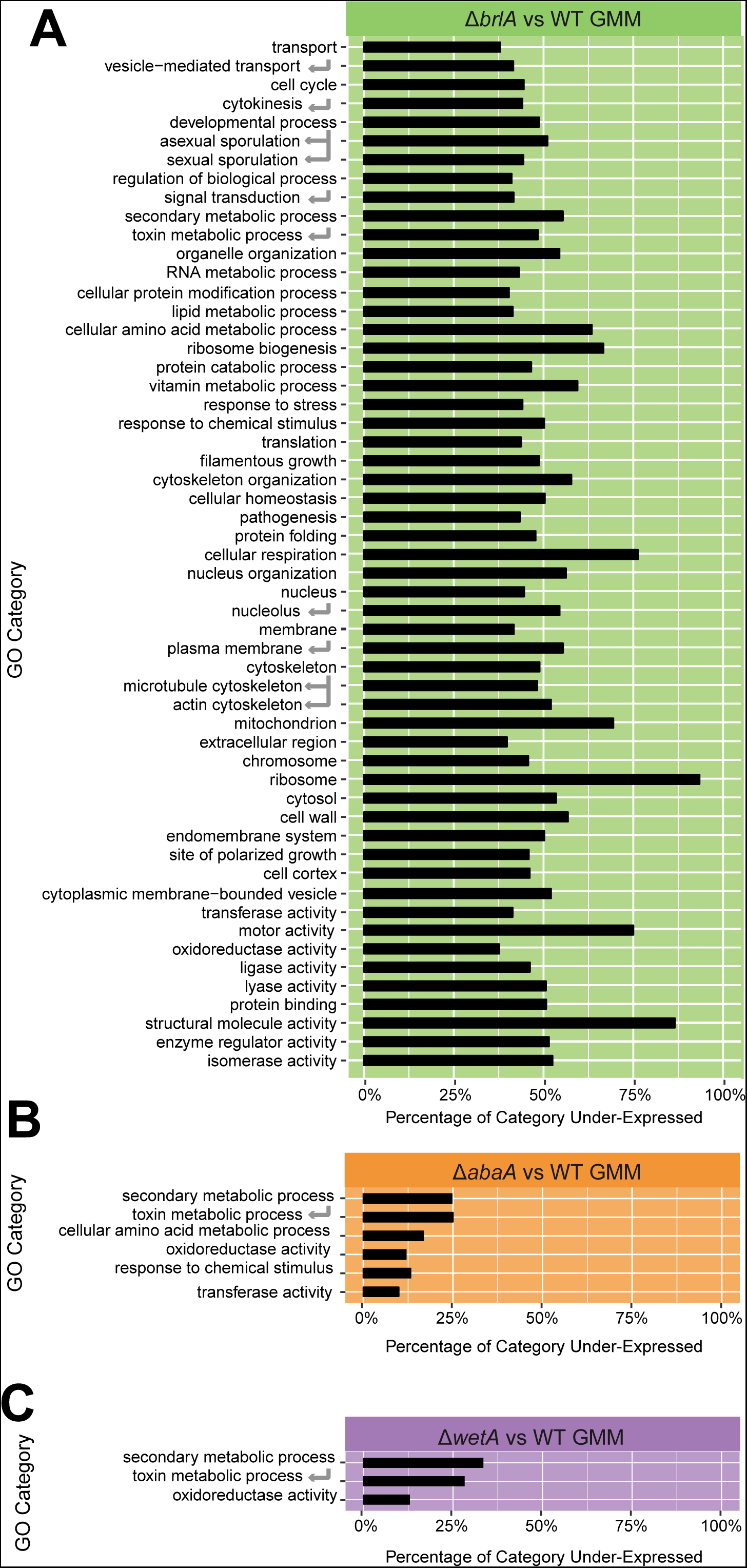
Genes under-expressed in Δ*brlA* are involved in a diverse set of cellular processes. Gene Ontology enrichment analysis for genes under-expressed in (A) Δ*brlA*, (B) Δ*abaA*, and (C) Δ*wetA* strains as compared to wild-type.

### BrlA is a key regulator of BGCs and SMs

To examine the relationship between regulation of asexual development and tissue-specific regulation of secondary metabolism, we examined the transcriptional responses of the 33 *A. fumigatus* characterized and predicted BGCs (**Table S3**) (29). Although only five characterized BGCs (DHN melanin, endocrocin, trypacidin, fumigaclavine and fumiquinazoline) are reported to be asexual development-specific, we found that 27 / 33 (82%) of BGCs are differentially expressed in one or more of the three comparisons examined (**Figure 2A**), suggesting a much broader governance of SM production by these transcriptional regulators of asexual development. Like the trend observed with genome wide transcriptional impact of these regulators, we find BrlA to be a major contributor to changes in BGC expression, regulating all but one of the differentially expressed BGCs (26 / 27; 96%), followed by WetA (15 / 27; 45%), and AbaA (11 / 27; 34%) (**Figure 2A**). Of the 27 differentially expressed BGCs, nine were regulated by all three transcriptional regulators, ten showed BrlA-specific regulation, one showed WetA-specific regulation, two showed joint regulation by both BrlA and AbaA, and five showed joint regulated by BrlA and WetA (**Figure 2B**). These results suggest that, unlike BrlA, WetA (with the exception of one BGC) and AbaA do not independently regulate their BGC targets.

**Figure 2.**
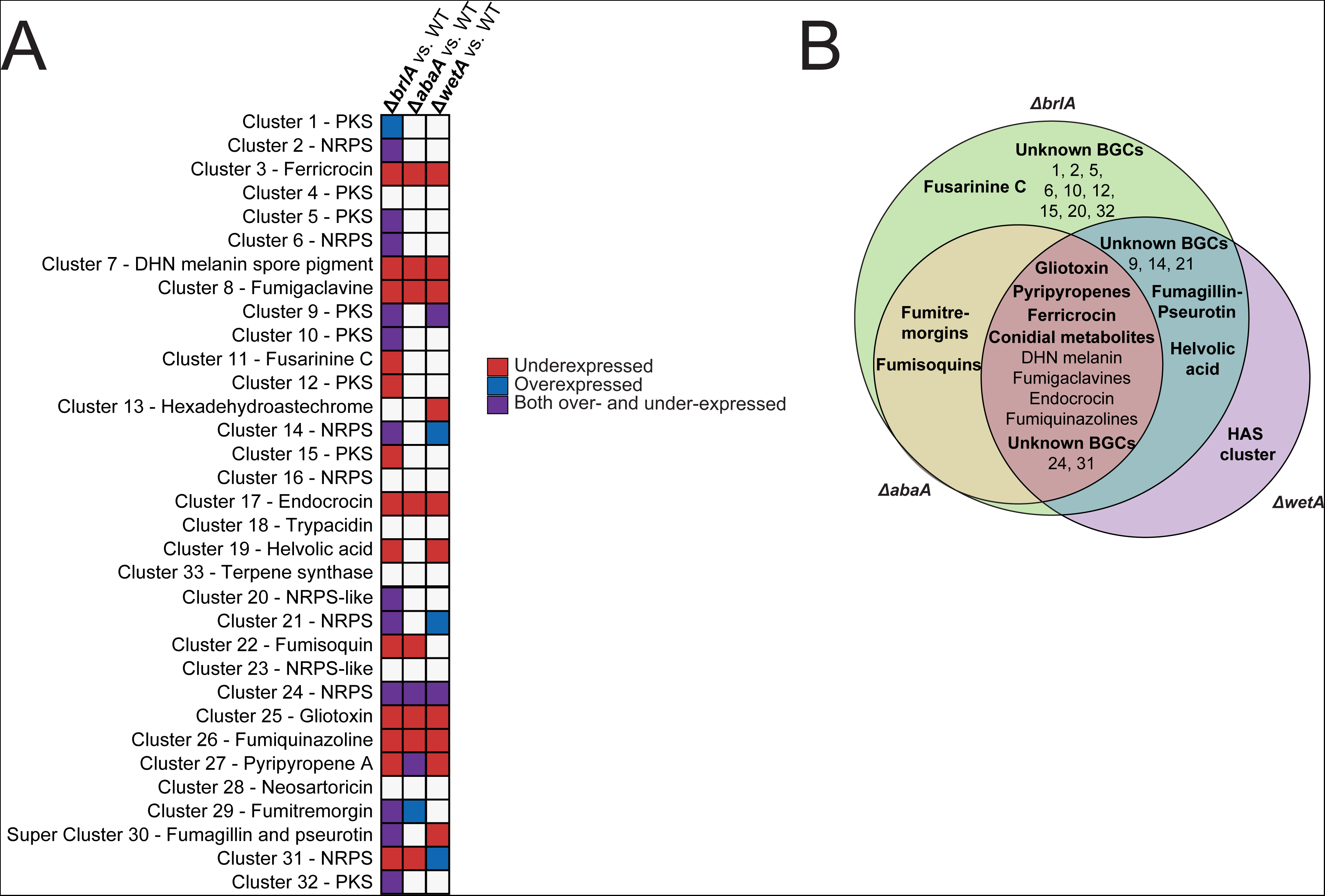
BrlA, AbaA, and WetA transcriptionally regulate many biosynthetic gene clusters (BGCs) involved in secondary metabolism. Expression of all secondary metabolic BGCs in *Aspergillus fumigatus* in all conditions tested. (A) Expression in Δ*brlA,* Δ*abaA,* and Δ*wetA*. (B) Overlap between BGCs under-expressed in Δ*brlA,* Δ*abaA,* and Δ*wetA*.

The nine BGCs that are jointly regulated by BrlA, AbaA, and WetA include the BGCs for ferricrocin, DHN melanin, fumigaclavines, endocrocin, gliotoxin, fumiquinazolines, and pyripyropenes, as well as the unknown non-ribosomal peptide synthetase (NRPS) cluster 24 and the unknown polyketide synthase (PKS) cluster 31 (**Figure 2**). Four of the five known asexual development-specific BGCs (endocrocin, fumigaclavine, fumiquinazoline, and DHN melanin) are regulated by all three developmental regulators (**Figure 2**); the only exception is the conidial PKS BGC for trypacidin, which appears to not be under the control of any of the developmental regulators in the conditions tested. However, the trypacidin pathway specific transcription factor gene *tpcE* (Afu4g14540), is positively regulated by BrlA (**Table S1**) and failure to observe *tpcC* (Afu4g14560; the polyketide synthase) regulation may be a reflection of observations that trypacidin is not produced at high levels until later time points (10). Aside from these four asexual development-specific BGCs, the other five BGCs that are jointly regulated by all three developmental regulators, include the gliotoxin BGC, the intracellular siderophore ferricrocin BGC, the meroterpene pyripyropene BGC, and BGCs 24 and 31 (**Figure 2**). Although previous results indicated the involvement of BrlA in regulating gliotoxin biosynthesis (16), it is not yet known whether this mycotoxin is present in asexual spores.

Joint regulation by BrlA, AbaA and WetA extends beyond the nine BGCs, and includes genes involved in sulfur/methionine metabolism (8 genes) and aromatic amino acid metabolism (4 genes) (**Table S1**) suggesting a connection between primary metabolism (e.g. substrate availability) and secondary metabolism. This is consistent with previous work linking methionine and tryptophan availability with natural product synthesis (19–21), as well as with evidence that *A. fumigatus* tryptophan metabolism mutants show altered secondary metabolite output (22). Specifically, many NRPS metabolites incorporate aromatic amino acids, such as tryptophan (e.g. fumiquinazoline) or phenylalanine (e.g. gliotoxin), in their carbon skeleton and gliotoxin itself impacts homeostasis of the methionine cycle (23, 24).

To examine whether the gene expression changes observed for these BGCs correlate with metabolite production, we performed SM profiling using the same fungal cultures as for the transcriptomic experiments (**Figure 3**). In the Δ*brlA* and Δ*abaA* mutant cultures, the metabolite profiles are consistent with the gene expression profiles of their corresponding BGCs. For example, production of ferricrocin, fumigaclavines, endocrocin, gliotoxin, fumiquinazolines, and pyripyropene A is completely abolished or significantly reduced in the Δ*brlA* mutant culture (**Figure 4**), mirroring the under-expression of their BGCs in the Δ*brlA* versus WT comparison (**Figure 2**). These compounds are also significantly reduced in the Δ*abaA* mutant culture and correlate with the gene expression patterns of their BGCs in the Δ*abaA* versus WT comparison, albeit to a lesser degree when compared to that observed in Δ*brlA* (**Figure 4; Figure 2**).

**Figure 3.**
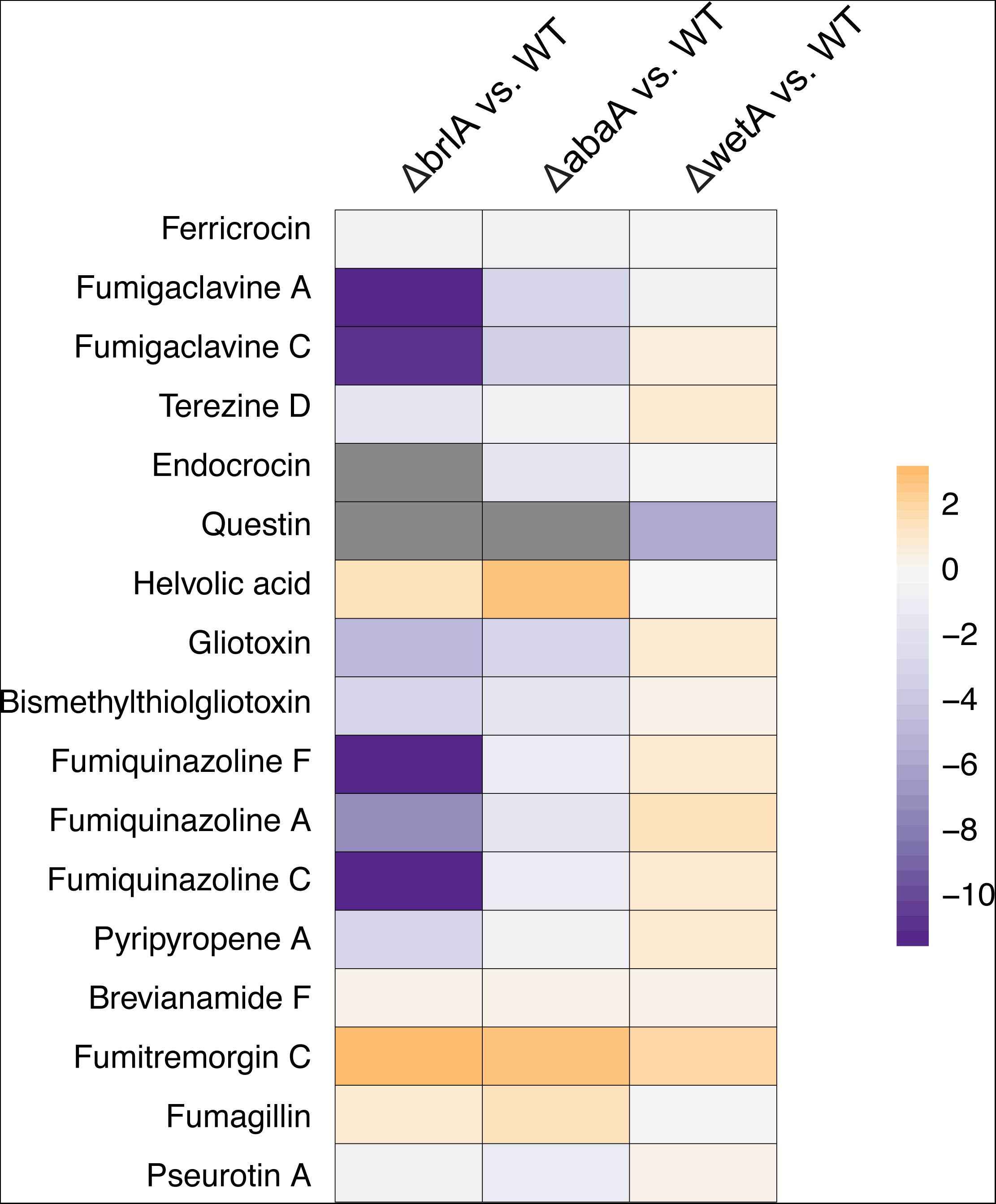
Secondary metabolites are produced at much lower concentrations in the *ΔbrlA* strain relative to wildtype. Summary of metabolite production in *ΔbrlA*, *ΔabaA*, and *ΔwetA* relative to the wildtype strain. Heat map colors represent log_2_ fold change in peak area intensity, and gray color indicates no metabolite was detected.

**Figure 4.**
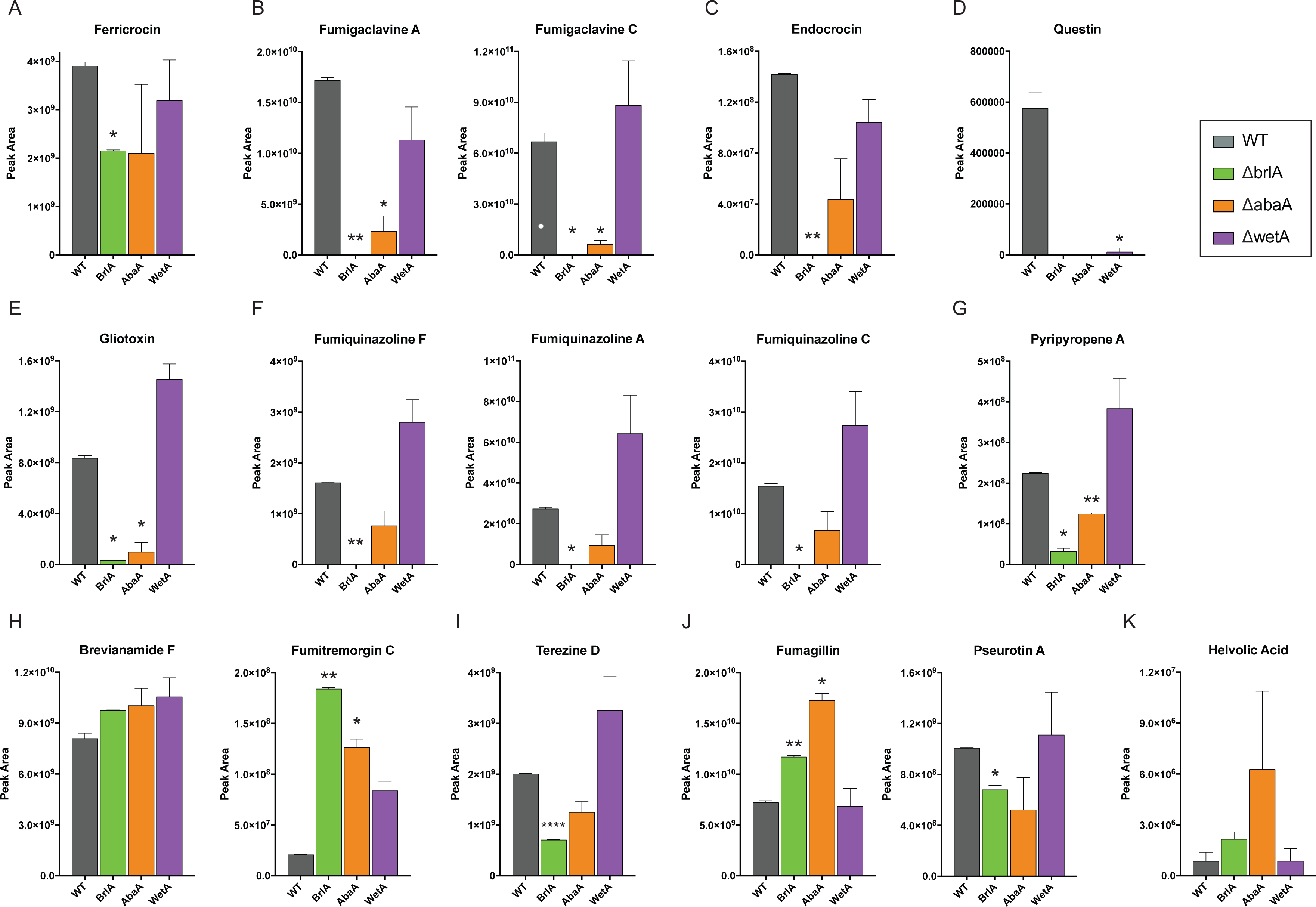
Concentrations of secondary metabolites produced by wild-type, Δ*brlA,* Δ*abaA,* and Δ*wetA* cultures. Peak area intensity of representative metabolites from differentially-expressed BGCs in *A. fumigatus*. Metabolite analysis was performed using the same two replicate samples from the RNA-seq experiment. Error bars depict standard deviation.

In contrast to Δ*brlA* and Δ*abaA*, the correlation between Δ*wetA* gene expression and metabolite profiles was much lower. For example, we observed a significant increase of several SMs, such as the fumigaclavines and endocrocin, in the Δ*wetA* mutant even though their corresponding BGCs are under-expressed in the Δ*wetA* versus WT comparison (**Figure 3; Figure 2**). Endocrocin is also produced as an early shunt product redundantly by the trypacidin BGC in the strain of *A. fumigatus* used in this study (10), and thus could be attributed to that BGC as well. Although we did not detect the final product trypacidin, minute amounts of the trypacidin precursor questin were produced in the wildtype fungus and, to much lesser degree, in the Δ*wetA* mutant (**Figure 4**). The high metabolite production levels in Δ*wetA* in spite of low gene expression levels could be attributed to the compromised cell wall of this developmental mutant (25) that resulted in increase in SM extraction efficiency compared to other test strains of the fungus.

Assessment of metabolites from both fungal tissue and growth supernatant (secreted) showed that a major fraction of extracted SMs, including all the known conidial-associated SMs (fumigaclavines, endocrocin, fumiquinazolines, and questin), the intracellular siderophore, ferricrocin, and pseurotin A accumulate in the fungal tissue, whereas that of other SMs, such as the fumitremorgins, terezine D, fumagillin, pyripyropene A, and helvolic acid, are secreted into the growth supernatant (**Figure S1, Table S4**). This may be a reflection of the chemical properties of these metabolites and their ability to diffuse or be actively released to the outside of the cell.

Ten of the 27 differentially regulated BGCs are under BrlA-specific control (**Figure 2B**). Except for the BGC of the extracellular siderophore fusarinine C, the SM products of the remaining nine BrlA-specific BGCs have yet to be characterized. Both BrlA and AbaA jointly govern expression of the fumisoquin BGC and the fumitremorgin BGC (**Figure 2B**). As the *A. fumigatus* Af293 strain used in this study is reported to harbor a point mutation in *ftmD* (Afu8g00200) that renders it incapable of producing the terminal product, fumitremorgin C (26), metabolomic analysis on the fumitremorgin BGC was performed using an early pathway precursor, brevianamide F. Total production of brevianamide F is significantly increased in cultures of all three transcriptional regulator mutants (**Figure 4**). Surprisingly, we also detect a significant amount of fumitremorgin C (as determined through m/zs and retention time matched to the fumitremorgin C standard) in the WT strain as well as – at even higher levels – in mutant cultures of all three developmental regulators, suggesting there may be compensation for this mutation (**Figure 4**). FtmD is an *O*-methyltransferase and it is possible that other *O*-methyltransferases in the genome may function at this step. On the other hand, Kato *et al.* (26) note that the mutated FtmD enzyme still functions and it is possible we observed fumitremorgin C production as we grew the fungus in a different condition than the one used in the previous report allowing for FtmD function.

Our gene expression data indicate that WetA positively regulates its sole specific target, the iron-coordinating hexadehydroastechrome (HAS) BGC. As metabolite detection of the iron coordination complex of HAS is challenging, we used the monomeric unit of this complex, terezine D, in our metabolite profiling of this BGC. In contrast to the gene expression data, which show that the BGC is under-expressed in Δ*wetA* versus WT, we observed that terezine D production is increased in the Δ*wetA* mutant (**Figure 4**). Even though the HAS BGC does not appear to be transcriptionally regulated by BrlA or AbaA, we still observed a decrease of terezine D production in both mutants (**Figure 4**). This could be related to other cellular processes as HAS is a tryptophan derived metabolite dependent on iron-homeostasis (27), with genes in both networks regulated by the BrlA cascade. BrlA and WetA jointly govern the helvolic acid BGC, the fumagillin/pseurotin supercluster, and three unknown BGCs (**Figure 2**). Compared to WT levels, production of both fumagillin and helvolic acid is increased in the Δ*brlA* mutant, unchanged in the Δ*wetA* mutant, and substantially increased in the Δ*abaA* mutant (**Figure 4**).

In summary, examination of transcriptional and metabolic profiles of Δ*brlA*, Δ*abaA*, Δ*wetA* and WT strains of *A. fumigatus* showed that several BGCs and SMs exhibit BrlA-specific regulation; in contrast, no BGCs or SMs were under AbaA-specific control and only one showed WetA-specific regulation. Furthermore, several additional BGCs and SMs appeared to be under the control of BrlA and WetA or AbaA or under the control of all three proteins (**Figure 2**). Given that strains lacking *brlA* do not enter asexual development, it is perhaps not surprising that both the gene expression and SM production of asexual development-specific BGCs, such as those for endocrocin and fumigaclavine, are under BrlA control (**Figure 2; Figure 4**). However, in addition to these spore-associated BGCs and SMs, BrlA also appears to regulate BGCs and SMs, such as helvolic acid and fumisoquin, which are not known to be associated with specialized developmental tissues but rather with vegetative growth, suggesting that BrlA regulation of secondary metabolism extends beyond asexual development.

### LaeA regulation of secondary metabolism is extensively mediated through BrlA

LaeA, a member of the fungal-specific velvet protein complex, is known to regulate secondary metabolism in many agriculturally and medically important filamentous fungi (28). Given the surprising global changes in BGC expression in the *ΔbrlA* mutant as well as the aberrant conidia phenotype previously observed in the *ΔlaeA* mutant (29), we further assessed the genetic relationship between these two global regulators and their governance on secondary metabolism. Global transcriptome comparison between the LaeA and BrlA regulons in *A. fumigatus* shows striking concordance in BGC regulation, with 13 / 16 of the LaeA-regulated BGCs as determined by microarray-based transcriptome analysis (13) also regulated by BrlA (**Table S5**). These include the BGCs responsible for the production of DHN melanin, fumigaclavines, endocrocin, helvolic acid, fumisoquins, gliotoxin, fumiquinazolines, fumitremorgins, fumagillin/pseurotin, and pyripyropenes, and three uncharacterized BGCs (cluster 24, a NRPS-based cluster upstream of the gliotoxin cluster, cluster 15, a PKS-based BGC and cluster 2, a nidulanin-like BGC) (**Table S5**). Unlike BrlA, which shows both positive and negative regulation of BGCs, LaeA strictly regulates BGCs in a positive manner in *A. fumigatus* at the time point assessed (13) (**Figure 5**). Interestingly, except for the fumitremorgin BGC, all of the jointly regulated BGCs are positively regulated by BrlA, further supporting a linked regulatory network between these two global regulators.

**Figure 5.**
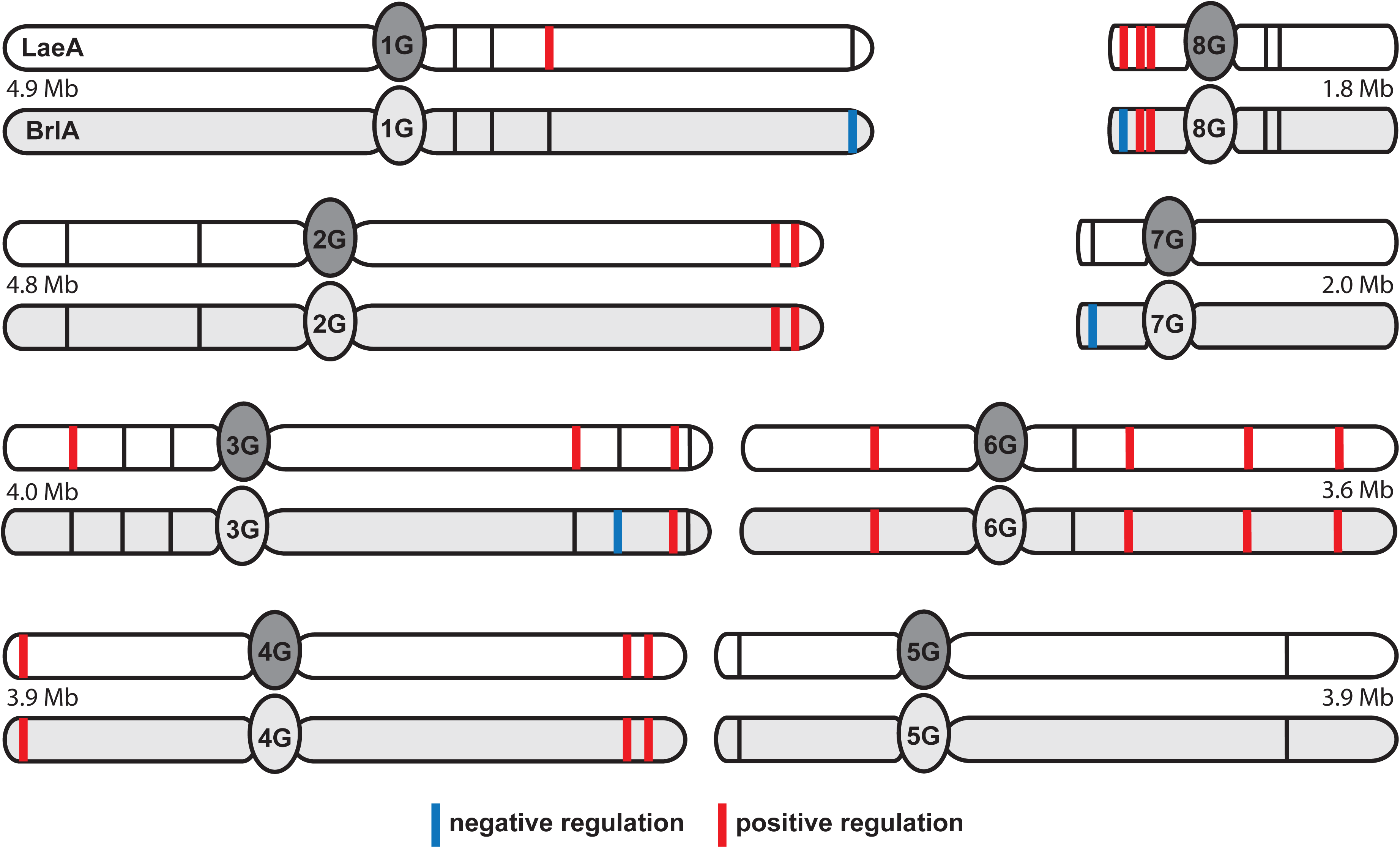
Chromosomal location of all biosynthetic gene clusters involved in secondary metabolism regulated by BrlA and by LaeA.

To further decipher the genetic relationship between the LaeA and BrlA transcriptional networks, we assessed the expression of these two genes in each respective deletion mutant. Northern analysis of both *brlA* and *laeA* expression in the *laeA* and *brlA* deletion mutants showed that whereas *laeA* expression is not significantly impacted in the Δ*brlA* strain (**Figure 6A**), *brlA* expression is significantly reduced in the *ΔlaeA* mutant (**Figure 6B**), in agreement with previous microarray and RNA-seq data (30, 31). Because LaeA loss is known to be involved in silencing of BGCs through chromatin remodeling (12) and a previous study in *A. nidulans* has shown that LaeA allows for SM expression by counteracting heterochromatin marks on BGC gene promoters, specifically reducing H3K9 methylation through heterochromatin protein-1, HepA (AN1905) (6), we suspected that the epistatic nature of LaeA regulation of *brlA* could be governed through modifications to the chromatin landscape within the *brlA* promoter. Indeed, chromatin immunoprecipitation (ChiP) examining histone modifications of the *brlA* promoter shows that although the histone H3 occupancy at the *brlA* promoter is unchanged between WT and *ΔlaeA*, there is a substantial decrease of a modification correlating with euchromatin (H3K4me3) in the *ΔlaeA* strain, while the heterochromatic mark H3K9-me3 is greatly enriched (**Figure 6C**). Thus, as with BGC regulation, it appears that LaeA epigenetically regulates *brlA* by impeding heterochromatin formation on the *brlA* promoter (**Figure 6**). Based on these results, we infer that LaeA regulation of secondary metabolism is significantly mediated through its epistatic effect on BrlA.

**Figure 6.**
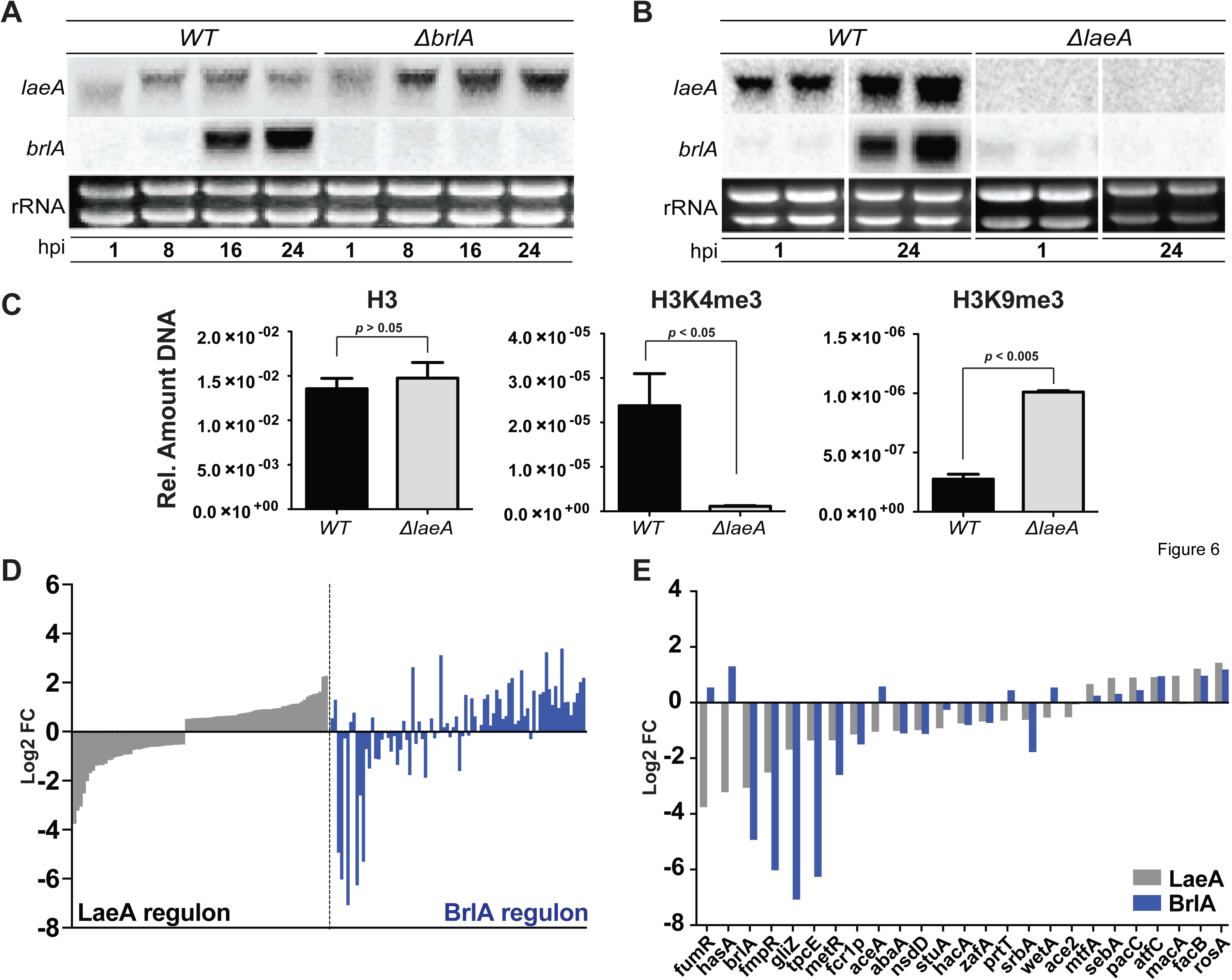
LaeA is epistatic to BrlA. (A) Northern analysis depicting expression of *brlA* and *laeA* in the *ΔbrlA* mutant compared to WT. (B) Northern analysis depicting expression of *brlA* and *laeA* in the Δ*brlA* mutant compared to WT. (C) Chromatin immunoprecipitation examining histone modifications of the brlA promoter. H3 depicts total Histone 3 occupancy. H3K4me3 depicts euchromatic mark. H3K9me3 depicts heterochromatic mark. Error bars depict standard deviation. (D) Predicted transcription factors in *A. fumigatus* with > 0.5 log_2_ fold change in the *ΔlaeA* mutant (left panel: grey). Log_2_ fold change of the same transcription factors in the *ΔbrlA* mutant (right panel: blue) (E) Log_2_ fold change of characterized transcription factors in the ΔlaeA (grey bars) and ΔbrlA (blue bars) mutants.

### Several transcriptional regulators of diverse cellular processes, including the SrbA-regulated hypoxia stress response, are also BrlA and LaeA-regulated

These findings piqued our interest on whether the LaeA-BrlA regulatory relay extends beyond secondary metabolism. To address this question, we compared the differential expression profiles of all *A. fumigatus* transcription factors (TFs) in both the Δ*laeA* versus WT comparison and in the Δ*brlA* versus WT comparison. We found that, similar to the observed overlap of differential expression profiles for BGCs (**Table S5**), many LaeA-regulated TFs are also regulated by BrlA (**Figure 6D**). A detailed assessment of functionally characterized TFs (**Figure 6E**) showed that both LaeA and BrlA positively regulate TFs found within BGCs (GliZ; Afu6g09630, FmpR/FapR; Afu6g03430, TpcE; Afu4G14550, FsqA; Afu03430) as well as a series of TFs involved in sulfur metabolism (MetR; Afu6g07530), sexual development (NsdD; Afu3g13870), asexual development (StuA; Afu2g07900, AbaA; Afu1g04830), the unfolded protein response (HacA; Afu3g04070), zinc response (ZafA; Afu1g10080), and the hypoxia response (SrbA; Afu2g01260) (**Figure 6D**). In addition, we also observe that both LaeA and BrlA negatively regulate a series of TFs involved in fungal morphogenesis (MtfA; Afu6g02690), virulence (SebA; Afu4g09080), pH signaling (PacC; Afu3g11970), acetate utilization (FacB; Afu1g13510), and repression of sexual development (RosA; Afu4g09710) (**Figure 6E**). It thus appears that a substantial part of the LaeA transcriptional cascade is moderated via BrlA.

To further examine a cellular process independent of secondary metabolism that is regulated by LaeA and BrlA, but not AbaA or WetA, we focused on the SrbA-regulated hypoxia stress response (32). Under-expressed genes in Δ*brlA* versus WT are enriched for categories associated with stress response and mitochondrial activity, including CELLULAR RESPIRATION, MITOCHONDRION, and RESPONSE TO STRESS (**Figure 1**). Among these genes are the hypoxia regulators *srbA* (Afu2g01260) and *srbB* (Afu4g03460) (**Table S1**). Both transcription factors contribute to virulence and are critical for regulation of iron uptake, heme biosynthesis and ergosterol synthesis in *A. fumigatus* (32). Previous work has determined that SrbA is a DNA-binding protein that binds upstream of 97 genes in *A. fumigatus* CEA10, 91 of which have orthologs in the Af293 strain used in this study (32). 69 / 91 (76%) of these genes are under-expressed in Δ*brlA* versus WT (**Table 2, Table S6**). In contrast, the percentages of SrbA-regulated genes were substantially smaller in both Δ*abaA* vs WT (14 / 91 genes or 15%) or Δ*wetA* vs WT (13 / 91 genes or 14%). Among the genes co-regulated by BrlA and SrbA are those in the ergosterol biosynthetic pathway, including the first enzyme in the pathway, Erg1 (Afu5g07780), both 14-α sterol demethylases (Erg11A/Cyp51A; Afu4g06890 and Erg11B/Cyp51B; Afu7g03740), Erg5 (Afu1g03950) and both C4-sterol methyl oxidases (Erg25A; Afu8g02440 and erg25B; Afu4g04820) (33, 34). The nitrate assimilation genes *niiA* (Afu1g12840) and *niaD* (Afu1g12830) are also regulated by BrlA and SrbA, linking sporulation and hypoxia to nitrate assimilation, observations noted in earlier studies (17, 35). Examination of the *ΔlaeA* transcriptional profile shows a near 100% identity of regulation of these genes (13). These findings largely replicate the working model for transcriptional regulation of the hypoxic response previously presented by Chung *et al.* (32), placing LaeA and BrlA as critical upstream regulators of this pathway.

**Table 2.**
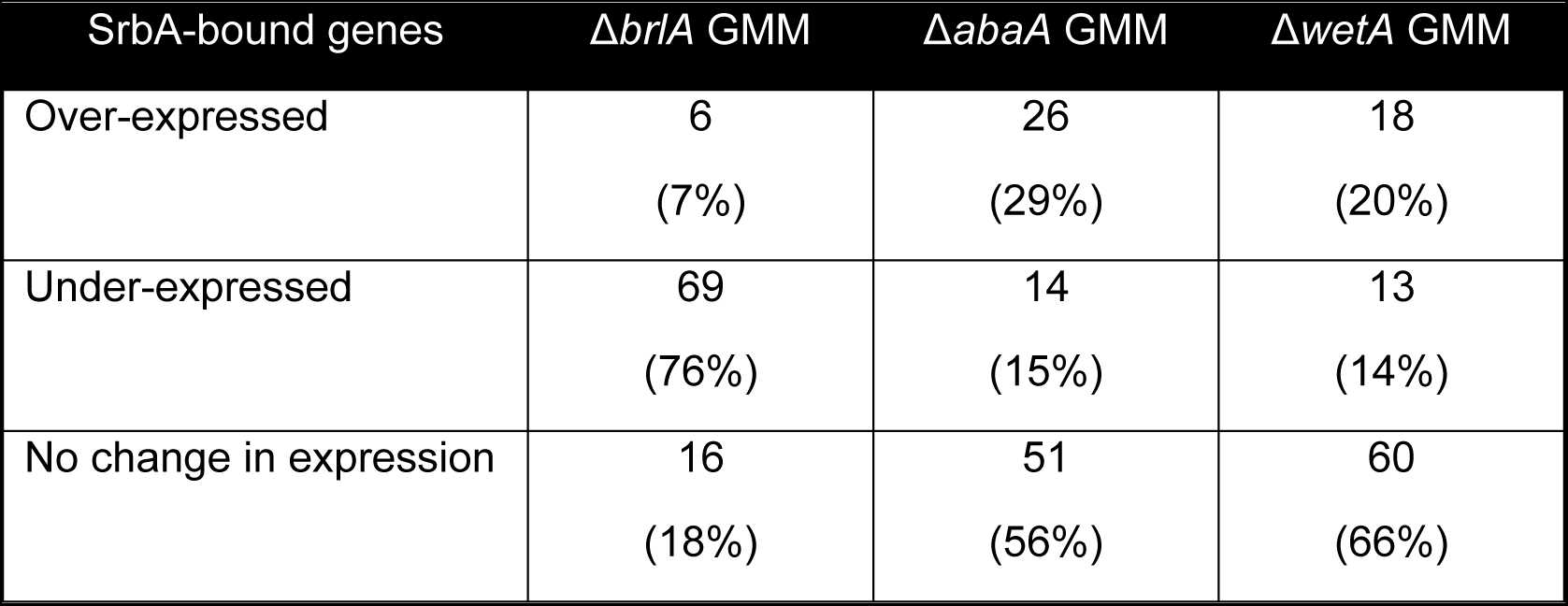
Expression of SrbA-bound genes in Δ*brlA,* Δ*abaA*, and Δ*wetA*.

### A cellular network regulating fungal secondary metabolism as well as diverse cellular processes

In filamentous fungi, SM production is coupled with the onset of asexual development. In *Aspergillus*, asexual development is governed by the central regulators BrlA, AbaA, and WetA, which are required for the early, middle, and late stages of asexual development, respectively. To investigate how regulation of asexual development is linked to the tissue-specific regulation of secondary metabolism, we examined the global transcriptomic and metabolomic profiles of Δ*brlA*, Δ*abaA*, Δ*wetA* and WT strains of *A. fumigatus.* We find a distinct role for BrlA in regulating both asexual development-specific and vegetative growth-specific secondary metabolism as well as diverse cellular processes, including the hypoxia stress response. Interestingly, BrlA’s involvement in SM regulation occurs in the context of the BrlA>AbaA>WetA cascade, whereas the protein’s involvement in the regulation of diverse cellular processes appears to be dissociated from AbaA and WetA. We further find that the BrlA transcriptional program is highly similar to the LaeA transcriptional program and elucidate the epigenetic and epistatic relationship of LaeA and *brlA* expression underlying the joint transcriptional profiles. Together, this work allows for a hierarchical framework of LaeA and BrlA function in fungal development (**Figure 7**).

**Figure 7.**
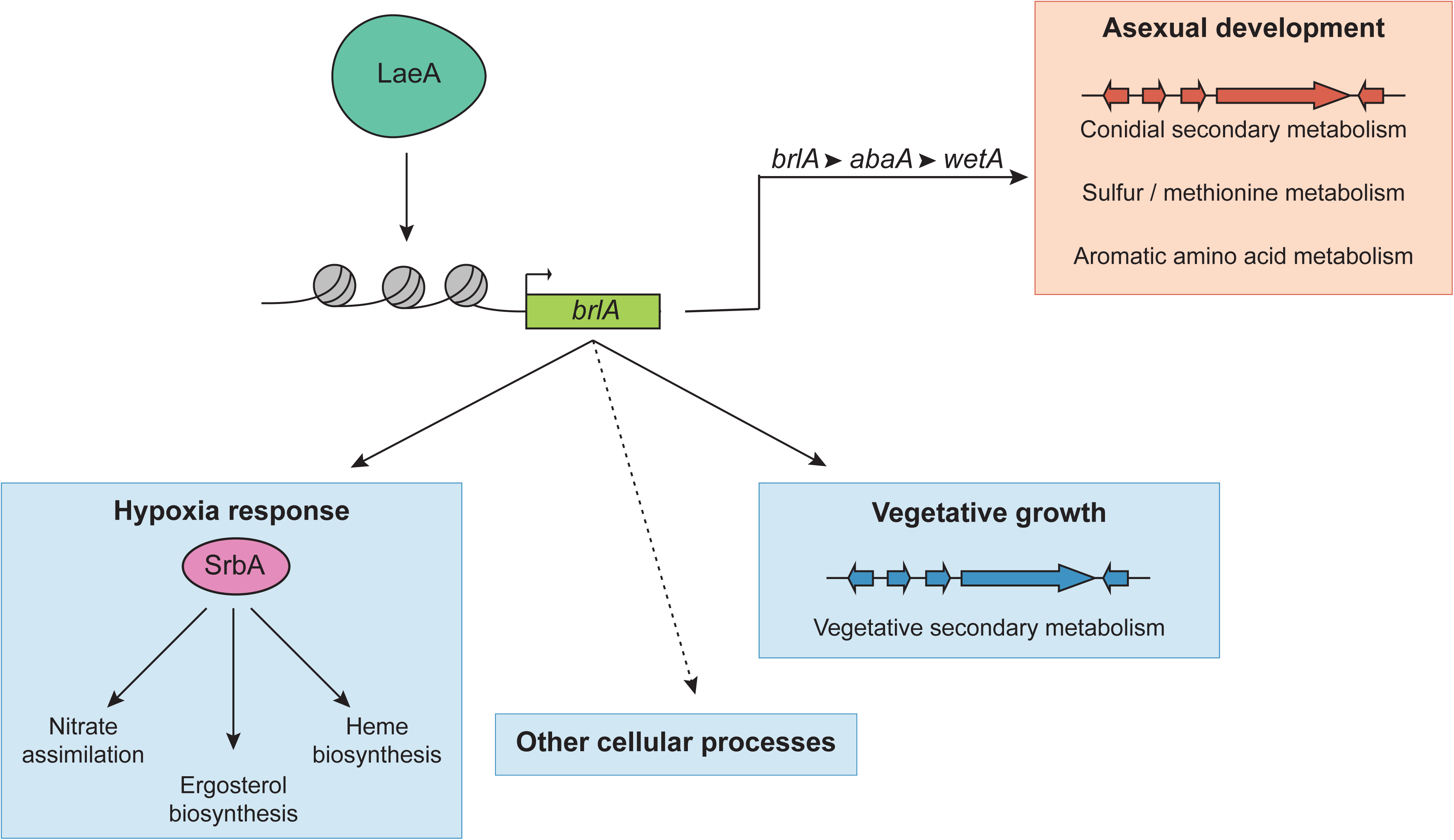
Model framework for the cellular network regulating fungal secondary metabolism and diverse cellular processes. Under our proposed model, the chromatin modifier LaeA, by epigenetically regulating the transcription factor BrlA, controls secondary metabolism in the context of both fungal vegetative growth and asexual development as well as additional cellular processes, such as the hypoxia response.

Both the set of BrlA>AbaA>WetA regulated genes and the set of BrlA / SrbA regulated genes show substantial overlap with the set of LaeA regulated genes (**Figure 6**; **Tables S4** and **S5**; 13, 31). Our finding that LaeA epigenetically regulates *brlA* expression provides a mechanistic explanation of these overlaps and alters our understanding of the role of LaeA in the regulation of secondary metabolism in the context of both fungal vegetative growth and asexual development as well as in the regulation of additional cellular processes. We propose that LaeA, perhaps as a member of the velvet protein complex, acts through epigenetic and epistatic regulation of key ‘cellular switches’, with BrlA representing one of these switches (**Figure 7**). BrlA is a known transcription factor that was first identified as a regulator of conidiophore development in *A. nidulans* (36, 37). BrlA has also been characterized in several *Aspergillus* and *Penicillium* species, and its regulatory functions have always associated with sporulation and frequently with secondary metabolism (16, 38–40). The regulatory elements in the *brlA* enhancer have been extensively characterized and are thought to include transcriptional complexes comprised of several regulatory proteins, including two velvet protein family proteins VosA and VelB (reviewed in Lee et al. 2016). It is possible that LaeA associates with one or more of these proposed positive acting transcriptional complexes to inhibit heterochromatic marks on the *brlA* promoter and allow for its activation.

The global nature of BGC regulation by BrlA was surprising and accounts for the majority of LaeA regulated SMs. Interestingly, of the nine characterized BGCs that our results suggest are identically regulated by both proteins, six do not contain a pathway-specific transcription factor and of the other three, the BGC-specific transcription factors (TpcE; Afu4g14540, FsqA; Afu6g03430, and GliZ; Afu6g09630, **Table S1**) are highly regulated by BrlA. Thus it appears that, minimally, these nine BGCs are induced by LaeA-mediated BrlA activation. However, not all BGCs were similarly regulated by LaeA and BrlA, suggesting that they may require LaeA activation through other or additional ‘cellular switches’, that they may be solely (positively or negatively) regulated by BrlA, or that they may be regulated through LaeA- and BrlA-independent cascades. Since both LaeA and BrlA are present in other fungal genera, including *Penicillium* and *Talaromyces*, and the fact that the secondary metabolites produced by organisms in these genera are distinct from those produced by *A. fumigatus*, it will be of future interest to address how conserved global molecular circuitry are rewired to control species-specific processes such as secondary metabolism (42).

Finally, our work shows that BrlA is the likely mediator of many of the known LaeA cellular cascades, including several associated with *A. fumigatus* virulence, substantially expanding the diversity of cellular processes that appear to be regulated by BrlA. For example, both proteins are critical for activation of members of the aromatic amino acid and sulfur/methionine pathways, which play a role in virulence of this pathogen (43, 44). We also find that BrlA is a key regulator of hypoxia regulated genes, likely through its regulation of SrbA and SrbB, the two key transcription factors critical for hypoxia adaptation in *A. fumigatus* (32, 45). SrbA is also important in azole resistance through its regulation of the ergosterol biosynthetic pathway (46), and our work uncovers a direct signaling pathway from LaeA to BrlA to SrbA/B to ergosterol gene expression which may reveal new avenues to study the expanding threat of antifungal resistance in *Aspergillus* species (47).

## Materials and Methods

### Fungal strains and growth conditions

All strains used in this study are listed in **Table 3**. Fungal strains are maintained in -80°C glycerol stocks and activated on glucose minimal media (GMM) at 37 °C (48). For RNA-seq analysis, 2.5 x10^6^ spores of WT, five confluent plates of Δ*brlA*, two confluent plates of Δ*abaA*, and 1 x 10^7^ spores of Δ*wetA* were inoculated into 500 mLs of liquid YPD (1% yeast extract, 2% peptone, 2% glucose) and grown in 250 rpm shaking condition for 24 hours at 37°C to synchronize development between strains. The size of the inoculum chosen was previously determined to provide comparable fungal mass after 24 hours incubation in the above condition. In a sterile environment, fungal mycelia were filtered through Miracloth (EMD Millipore) and thoroughly washed in PBS to remove residual YPD. Equal amounts of mycelia were transferred into three flasks containing 250 mLs of liquid GMM and incubated at 30 °C in 250 rpm shaking condition to induce development and conidiation (29). Approximately equal amounts of mycelia were removed from all fungal strains at 48 hours post induction, flash frozen in liquid nitrogen, lyophilized, and stored at -80°C until being used for RNA extraction and downstream RNA sequencing. For metabolomics analysis, all fungal cultures were left to incubate and total culture (growth supernatant and fungal mycelia) were harvested at 96 hours post induction, frozen in -80°C until ready to be extracted and analyzed.

**Table 3.** Strains used in this study.

| Strain | Identifier | Genotype | Source |
| --- | --- | --- | --- |
| Af293 | WT | Wild type | (61) |
| TJW54.1 | $\Delta laeA$ | $\Delta laeA::parapyrG$; *pyrG1* | (29) |
| $\Delta AfbrlA7$ | $\Delta brlA$ | $\Delta brlA::fumipyrG$; *pyrG1* | (62) |
| TSGa17 | $\Delta abaA$ | $\Delta abaA::fumipyrG$; *pyrG1* | (25) |
| TSGw4 | $\Delta wetA$ | $\Delta wetA::fumipyrG$; *pyrG1* | (25) |

### RNA isolation and sequencing

Total RNA was extracted with QIAzol^®^ reagent (QIAGEN^®^) from freeze-dried mycelia harvested at 48 hours post induction of asexual development following manufacturer’s protocol and further purified using silica membrane spin columns from RNeasy^®^Plant mini kit (QIAGEN^®^). Total RNA was subjected to DNaseI digestion to further remove genomic DNA contamination. RNA-seq libraries were constructed and sequenced at the Genomic Services Lab of Hudson Alpha (Huntsville, Alabama) using 50bp Illumina paired-end stranded reads. Libraries were constructed with the Illumina TruSeq Stranded mRNA Library Prep Kit (Illumina) and sequenced on an Illumina HiSeq 2500 sequencer. Two biological replicates were generated for each strain sequenced and 28 – 53 million reads were generated for each library. All short read sequences are available in the NCBI Short Read Archive under the BioProject PRJNA396210.

### Differential gene expression analysis

Raw RNA-seq reads were trimmed of low-quality reads and adapter sequences using Trimmomatic with the suggested parameters for paired-end read trimming (49). After read trimming, all samples contained between 19-49 million read pairs, with the average sample containing 28 million reads. Trimmed reads were aligned to the *A. fumigatus* Af293 version s03_m04_r11 genome from the *Aspergillus* Genome Database (50, 51). Read alignment was performed with Tophat2 using the reference gene annotation to guide alignment and without attempting to detect novel transcripts (parameter –no-novel-juncs) (52). Reads aligning to each gene were counted using HTSeq-count with the union mode (53). Differential expression was determined using the DESeq2 software (54). Genes were considered differentially expressed if their Benjamani-Hochberg adjusted p-value was less than 0.1.

### Functional enrichment analysis

Functional category enrichment was determined for differentially expressed genes in all conditions tested using the Cytoscape plugin BiNGO (55, 56). To allow for a high-level view of the types of differentially expressed gene sets, the Aspergillus GOSlim v1.2 term subset was used (57). The Benjamini-Hochberg multiple testing correction was applied and functional categories were considered significantly enriched if the adjusted p-value was less than 0.05.

### Gene cluster expression

*A. fumigatus* BGCs were taken from a combination of computationally predicted and experimentally characterized gene clusters involved in secondary metabolism (18, 58). A list of all BGCs used in this study is available in **Table S2.** BGCs were designated differentially expressed if half or more of the genes in the BGC were differentially expressed. BGCs were designated over-expressed if half or more of the genes in the BGC were over-expressed or were designated under-expressed if half or more of the genes in the BGC were under-expressed. BGCs where half or more of the genes in the BGC were differentially expressed but did not have half or more genes being either over-expressed or under-expressed were designated mixed expression.

### Metabolomics analysis

#### Metabolite extraction

Total culture (growth supernatant and fungal mycelia) were obtained 96 hours post induction of asexual development. Growth supernatant and fungal mycelia were separated via filtration through Miracloth (EMD Millipore). Prior to extraction of the growth supernatant, residual mycelial debris were pelleted via centrifugation and 5 mLs of the growth supernatant were subjected to solid phase extraction (SPE) using EVOLUTE^®^ ABN SPE columns (Biotage) following manufacturer’s protocol. The fungal mycelia were washed thoroughly with PBS to remove residual growth supernatant and extracted using ethyl acetate/dichloromethane/methanol (3:2:1; v/v/v) with 1% (v/v) formic acid (59) coupled with incubation in a sonicator water for one hour. Both the growth supernatant and fungal mycelial crude extracts were evaporated to dryness using a Thermo Scientific Savant SC250 vacuum concentrator and stored at -20°C until ready for ultra-high performance liquid chromatography-mass spectrometry (UHPLC-MS) analysis.

#### UHPLC-MS analysis: equipment overview and analytical methods

High resolution UHPLC-MS was performed on a Thermo Scientific Vanquish™UHPLC system coupled to a Thermo Scientific Q Exactive hybrid quadrupole Orbitrap™MS. The system was operated in both electrospray positive (ESI^+^) and electrospray negative (ESI^-^) modes with ion voltages set at 3.5 kV in both modes.

Crude extracts were reconstituted in 0.5 mL of 50% (v/v) acetonitrile + 0.1% (v/v) formic acid and syringe filtered through the 0.2 μM PTFE filter to remove insoluble materials. 10 μL were injected into the UHPLC-MS system, separated using an Agilent Zorbax Eclipse XDB-C18 column (2.1 x 150 mm, 1.8 μM particle diameter), and ran using 0.05% formic acid in acetonitrile as the organic phase and 0.05% formic acid in water as the aqueous phase at a flow rate of 0.2 mL/min. The solvent gradient starts at 20% organic for 2 mins, followed by a linear increase to 60% organic over 10 minutes, a linear increase to 100% organic over 1 min, and a final holding at 100% organic for 5 mins totaling to 18 minutes of runtime and data collection. The XDB-C18 column was equilibrated at 20% organic for 5 mins in between each sample injection throughout the entire sequence.

Purified standards were used to validate compounds analyzed in this study. Standards used were either commercially purchased or kindly gifted by other investigators as described below: helvolic acid (21580; Cayman Chemical Company), gliotoxin (G9893; Sigma-Aldrich), brevianamide F (HY-100385; MedChem Express), fumagillin (11332, Cayman Chemical Company), pseurotin A (14441, Cayman Chemical Company), fumitremorgin C (11030; Cayman Chemical Company), pyripyropene A (11896, Cayman Chemical Company), terezine D (purified and gifted by the Schroeder lab at Cornell University), endocrocin and questin (purified and gifted by the Wang lab at University of Southern California), fumiquinazolines F and A (purified and gifted by the Walsh lab at Harvard Medical School), trypacidin (purified and gifted from the Puel lab at French National Institute for Agricultural Research – Toulouse), fumigaclavine A (SC-203051; Santa Cruz Biotechnology). Standards for ferricrocin and fumigaclavine C were unavailable and thus compound abundance were inferred from calculated *m/z*.

Data visualization, peak alignment, analysis of full scan UHPLC-MS data, ion extraction, and metabolite quantitation were performed using Xcalibur™(Thermo Scientific) and MAVEN (Melamud et al, 2010). Ionization mode was chosen for each compound based on optimal peak profiles of their respective standards as assessed in both ESI^+^ and ESI^-^ modes. Both total ion chromatograms (TICs) and extracted ion chromatograms (EICs) were generated in GraphPad Prism 7 (GraphPad Software) using coordinate data of peak intensity (y) vs retention time (x) obtained from MAVEN. The area below the peak that corresponds to each compound was used to generate the table for metabolite quantitation in GraphPad Prism 7 (GraphPad Software).

### Chromatin immunoprecipitation and real time qPCR analysis

50 milliliter cultures of liquid GMM were inoculated with 1 x 10^6^ spores per ml and incubated at 250 rpm and 37 °C for 24 hours under light. Triplicate cultures were performed for each strain. Chromatin immunoprecipitation was carried out as described previously (60). Antibodies used for ChIP were: rabbit polyclonal to histone H3 acetyl K9, Abcam, ab10812, rabbit polyclonal to histone H3 trimethyl K4, Upstate, 07-473, rabbit polyclonal to histone H3 acetyl K9, Abcam, ab8898, and rabbit polyclonal to C-terminus histone H3 antibody, ab1791. Two micrograms of antibody were used per reaction of 200 mg total protein. Amplification and detection of precipitated DNA in real-time qPCR was performed with iQ™ SYBR^®^ Green Supermix #170-8880 (Bio-Rad, Cat#170-8880) following the manufacturer’s instructions using primers AF brlA(p) F qPCR (CGTACGGGTGTAAGTCTGATC) and AF brlA(p) R qPCR (CTCTGTATCTTCTAGTTCAATGG). Relative amounts of DNA were calculated by dividing the immunoprecipitated DNA by the input DNA. Each PCR reaction was replicated. To normalize the amount of DNA precipitated with histone H3-acetyl K9 and H3-trimethyl K4, the quantities from precipitation with these antibodies was divided by the previously calculated ratio of the anti-C-terminus histone H3 precipitation to input DNA.

## Acknowledgements

A.L.L. was supported by the U.S. National Library of Medicine training grant 2T15LM007450. This work was supported in part by the National Science Foundation (DEB-1442113 to A.R.), by the National Institutes of Health (R01 AI065728-01 to N. P. K. and T32 GM07133 and NRSA AI55397 to A. A. S.). This work was conducted in part using the resources of the Advanced Computing Center for Research and Education at Vanderbilt University (http://www.accre.vanderbilt.edu/).

## Supplementary Material

**Figure S1. Concentrations of secondary metabolites produced by wild-type, Δ*brlA,* Δ*abaA,* and Δ*wetA* cultures and located in mycelia and supernatant.** Peak area intensity of representative metabolites from differentially-expressed BGCs in *A. fumigatus*. Metabolite analysis was performed using the same two replicate samples from the RNAseq experiment. Error bars depict standard deviation. Statistics are located in Table S4

**Table S1.** Differential gene expression of all strains.

**Table S2.** All Gene Ontology enrichment results.

**Table S3.** All secondary metabolic gene clusters in *Aspergillus fumigatus*.

**Table S4.** T-tests for significant differences between supernatant and mycelial secondary metabolites shown in Figure S1.

**Table S5**. Overlap in LaeA and BrlA regulated clusters.

**Table S6**. Shared targets of SrbA and BrlA.

